# Cell morphology QTL reveal gene by environment interactions in a genetically diverse cell population

**DOI:** 10.1101/2023.11.18.567597

**Authors:** Callan O’Connor, Gregory R. Keele, Whitney Martin, Timothy Stodola, Daniel Gatti, Brian R. Hoffman, Ron Korstanje, Gary A. Churchill, Laura G. Reinholdt

## Abstract

Genetically heterogenous cell lines from laboratory mice are promising tools for population-based screening as they offer power for genetic mapping, and potentially, predictive value for *in vivo* experimentation in genetically matched individuals. To explore this further, we derived a panel of fibroblast lines from a genetic reference population of laboratory mice (the Diversity Outbred, DO). We then used high-content imaging to capture hundreds of cell morphology traits in cells exposed to the oxidative stress-inducing arsenic metabolite monomethylarsonous acid (MMA^III^). We employed dose-response modeling to capture latent parameters of response and we then used these parameters to identify several hundred cell morphology quantitative trait loci (cmQTL). Response cmQTL encompass genes with established associations with cellular responses to arsenic exposure, including *Abcc4* and *Txnrd1*, as well as novel gene candidates like *Xrcc2*. Moreover, baseline trait cmQTL highlight the influence of natural variation on fundamental aspects of nuclear morphology. We show that the natural variants influencing response include both coding and non-coding variation, and that cmQTL haplotypes can be used to predict response in orthogonal cell lines. Our study sheds light on the major molecular initiating events of oxidative stress that are under genetic regulation, including the NRF2-mediated antioxidant response, cellular detoxification pathways, DNA damage repair response, and cell death trajectories.

## Introduction

Cell morphology has served as a useful phenotype for understanding how genetic factors regulate the state of metazoan cells, ranging from yeast to human induced pluripotent stem cells (iPSCs) ^1,2^. Recent advances in microscopy-based, high-content cellular screening (HCS) have made it cost-effective to analyze cellular phenotypes at scale ^3–7^. When coupled with machine learning techniques, these technologies enable precise measurements of cellular and sub-cellular morphological traits, which have long been observed in the context of development and disease ^8–11^.

We and others previously characterized the genetic architecture of ground-state pluripotency and differentiation propensity in genetically diverse mouse embryonic stem cells (mESCs). This work demonstrated that -omics traits like gene expression, chromatin accessibility, and protein levels in genetically diverse cells, especially when combined (multi-omics), provide molecular readouts that can be used to identify the genetic factors regulating cell state ^12–15^. The correlation of cell morphology traits to these underlying -omics traits offers the potential to quantitatively analyze and delineate how cells respond to genetic and environmental perturbations ^16–18^. However, multi-omic approaches like these can be expensive, particularly in the context of population-level screens of cell state across many environmental perturbations. Moreover, the utility of cell morphology traits derived from HCS for genetic analysis has not been fully explored, especially in laboratory mouse cells.

In this study, we used cell morphology traits from HCS for genetic analysis of cellular response during acute arsenic exposure. Arsenic is a known carcinogen and a widespread contaminant of groundwater, exposing up to estimated 220 million people worldwide ^19^. Ingested inorganic arsenic is metabolized through methylation and reducing reactions that generate metabolites including monomethylarsonic acid (MMA^V^), monomethylarsonous acid (MMA^III^), dimethylarsinic acid (DMA^V^), and dimethylarsinous acid (DMA^III^) ^20–22^. These arsenic metabolites have unique toxicological profiles and urinary ratios that favor the more toxic forms have been linked to disease ^23,24^. At the cellular level, arsenic exposure induces oxidative stress, DNA damage, and cytotoxicity to varying degrees depending on the metabolites present, the tissue type, and genetic background of the exposed individual. These are the key events that lead to adverse outcomes including cancer or impaired reproduction / development at the population level. Interindividual variation in urinary metabolite ratios from populations exposed to high levels of arsenic have been used in genetic association mapping to identify variants associated with adverse outcomes in sensitive individuals. These studies revealed genes and variants that regulate arsenic metabolism, as well as oxidative stress response and DNA damage repair ^25–41^. In laboratory mice, the metabolite MMA^III^ causes DNA damage through oxidative stress and induces tumor development in the kidney ^42,43^. Given the substantial body of genetic association data for arsenic and our interest in kidney pathophysiology, we sought to evaluate a population-based cellular model and to employ cell morphology traits to access gene by environment interactions for the metabolite MMA^III^.

Genetically diverse laboratory mouse resource populations are powerful experimental tools for genetic analysis and they are well established in the study of gene by environment interactions *in vivo* ^44,45^. Cell lines from these genetic reference populations offer a new approach methodology wherein genetic screens can be performed ‘in a dish’ to identify haplotypes that confer sensitivity and resilience. Approaches such as these have the potential to reduce the scale of animal studies where informative molecular and/or cellular phenotypes exist. We created a diverse panel of primary fibroblast cell lines from the Diversity Outbred (DO) mouse population^46^. DO mice are outbred animals descended from eight inbred mouse strains: A/J (AJ), C57BL/6J (B6), 129S1/SvImJ (129), NOD/ShiLtJ (NOD), NZO/HILtJ (NZO), CAST/EiJ (CAST), PWK/PhJ (PWK), and WSB/EiJ (WSB). These inbred strains represent three sub-species of *Mus musculus* and thus possess far more genetic variation than traditional mouse crosses, capturing roughly 45 million segregating single nucleotide polymorphisms (SNPs) ^46,47^.

Using a high content screening (HCS) technique similar to Cell Painting ^3^, we show that high-dimensional cell morphology phenotypes can be summarized through dose-response modeling to capture latent features that reflect changes in cell state during an acute, arsenic-induced oxidative stress response. We show that these cell state changes vary across genetically diverse cells, revealing both sensitive and resilient individuals to MMA^III^-induced cell morphology changes. Using quantitative trait mapping (QTL), we found 854 cell morphology QTL (cmQTL; LOD score > 7.5), which are the genetic loci that regulate the cellular response to arsenical exposure. Additionally, we show that the cmQTL effects are both reproducible and predictive of arsenic sensitivity. At the gene and pathway level, many cmQTL recapitulate genetic associations that have been previously found in human population studies, demonstrating the translational utility of our population-based cellular model. We highlight the roles of *Xrcc2 and Txnrd1* alleles that modulate MMA^III^-induced cellular death, and we provide new associations for a host of candidate genes that interact with MMA^III^.

## Results

Cell morphology is influenced by genetic variation and environmental factors including chemical exposures ^2^. Therefore we sought to use morphological traits to quantify the key cellular events that occur during arsenic exposure, and to identify the genetic determinants of cellular sensitivity through a forward genetic screen. We established a population-based cellular model by deriving a panel of tail tip fibroblast lines from the Diversity Outbred (DO) mouse population (n = 600) (**Fig. 1A,1B**). Tail tip fibroblast cultures can be readily established through minimally invasive techniques, they are adherent, and they can be easily maintained for many passages depending on the age of the donor. Though heterogeneous and tissue specific, fibroblasts are one of the most widespread cell types found in mammals. To observe effects of acute arsenic exposure, we treated 226 of these DO fibroblast lines with eight increasing concentrations of monomethylarsonous acid (MMA^III^) across 76 randomized 96-well plates ^48^. MMA^III^ is a highly toxic arsenic intermediate that induces oxidative stress associated DNA damage in exposed tissues ^49^ (**Fig. 1A**). Based on the genetic architecture of the DO population, we expected this number of individual cell lines would allow us to detect QTL explaining >20% of the phenotypic variance with 90% power ^50^. To quantify changes in cell morphology associated with oxidative stress and genotoxicity, we used cell stains to label nuclei (Hoechst 33342) and mitochondria (MitoTracker Deep Red), and we used indirect immunolabeling to quantify DNA damage repair (γH2AX) (**Fig. 1C**). We captured 180,255 images and performed image analysis using Harmony 4.9 to extract 673 image-based, morphological phenotypes from 2,721,560 cells (**Fig. 1B)**.

**Figure 1:**
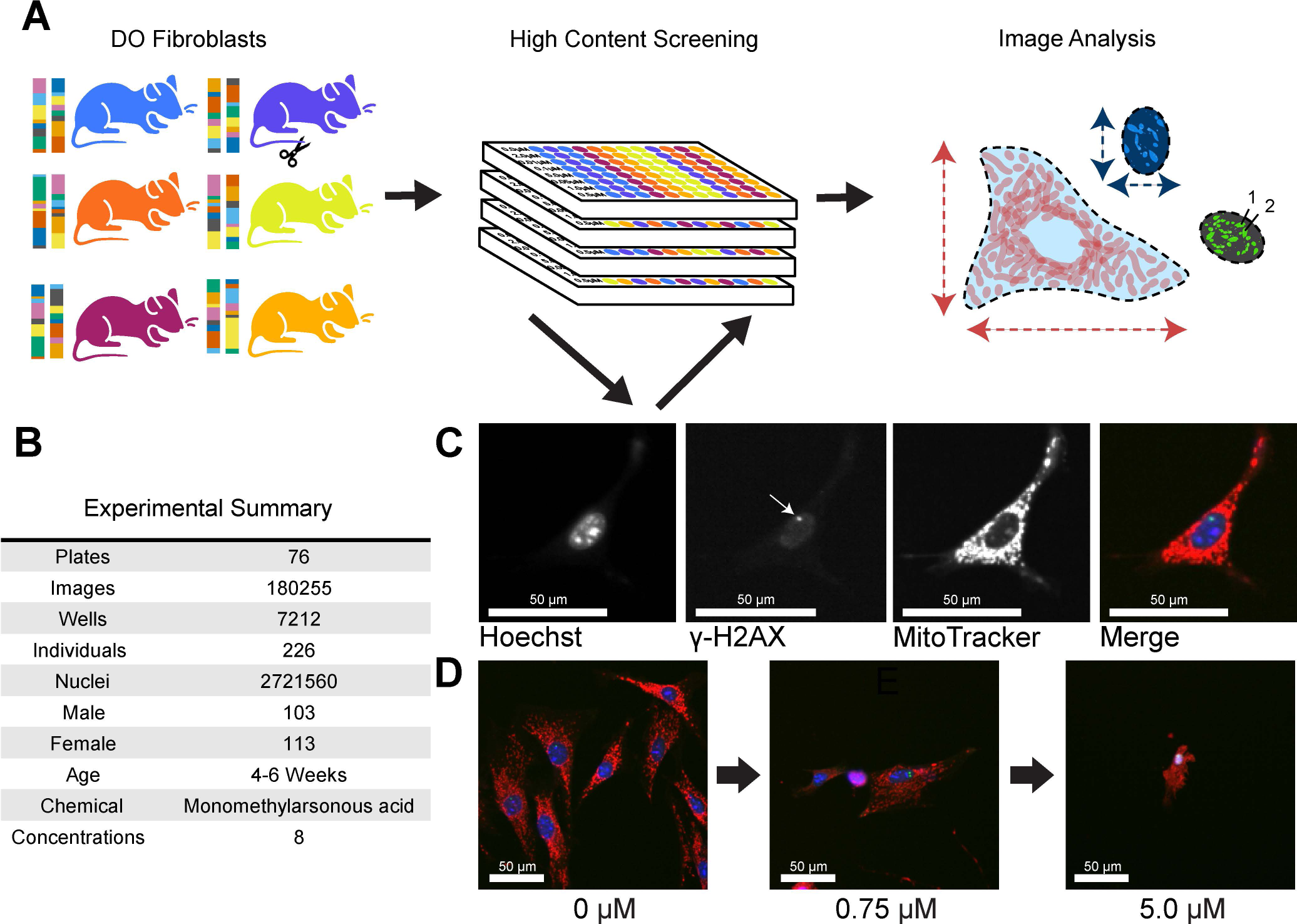
HCS of MMA^III^-exposed DO Fibroblasts. (A) 600+ primary fibroblasts were derived from Diversity Outbred (DO) mice aged 4-6 weeks. 226 DO fibroblast lines were exposed to 8 concentrations of MMA^III^ (0 µM, 0.01 µM, 0.1 µM, 0.75 µM, 1.0 µM, 1.25 µM, 2.0 µM, and 5.0 µM). Cell lines were semi-randomly seeded into 96-well plates (4 columns spanning two plates, see Supplementals for more information). Image analysis was performed at the whole well level and summarized across concentrations using dose response modeling. (B) Table with experimental summary (C) Example images showing fibroblasts labeled with MitoTracker Deep Red, Hoechst 33342, an anti-gamma γH2AX antibody with a Alexafluor 488 donkey anti-rabbit secondary, and the merged image. Plates were imaged using an Operetta High Content Imager (PerkinElmer) at 20X. D) Example merged images showing a fibroblasts’ morphology across three representative doses of MMAIII (0 µM, 0.75 µM, 5.0 µM).

### Sources of variation in cell morphology traits

To assess the main drivers of variation in these data, we performed principal components analysis. The first principal component, accounting for 41.5% of the observed variation across all traits was correlated with MMA^III^ concentration, and there was a clear dose-dependent effect (**Fig. 2A**). Following Matthew et al. ^2^, we performed a decomposition of the sources of variation contributing to each trait by fitting a random effects linear model with terms for inter-plate effects (‘plate’), batch effects (12 samples per ‘run’), MMA^III^ concentration (‘concentration’), DO donor (‘individual’), and the sex of cell donor (sex) (**Fig. 2B**). Among these factors, arsenic ‘concentration’ explained the most variation, followed by ‘individual’ or donor genetic background. While we randomized DO cell lines by column and MMA^III^ concentrations by row within a plate, we observed a common HCS finding that inter-plate and inter-run effects also influence variance in measured cellular features (**Fig. 2B**). Depending on the trait, ‘individual’ explained ∼0-40% of the variance with an average of 10%, suggesting that a subset of these traits (those with >20%) would provide sufficient signal for genetic mapping based on the size and architecture of our DO cell population^50^.

**Figure 2:**
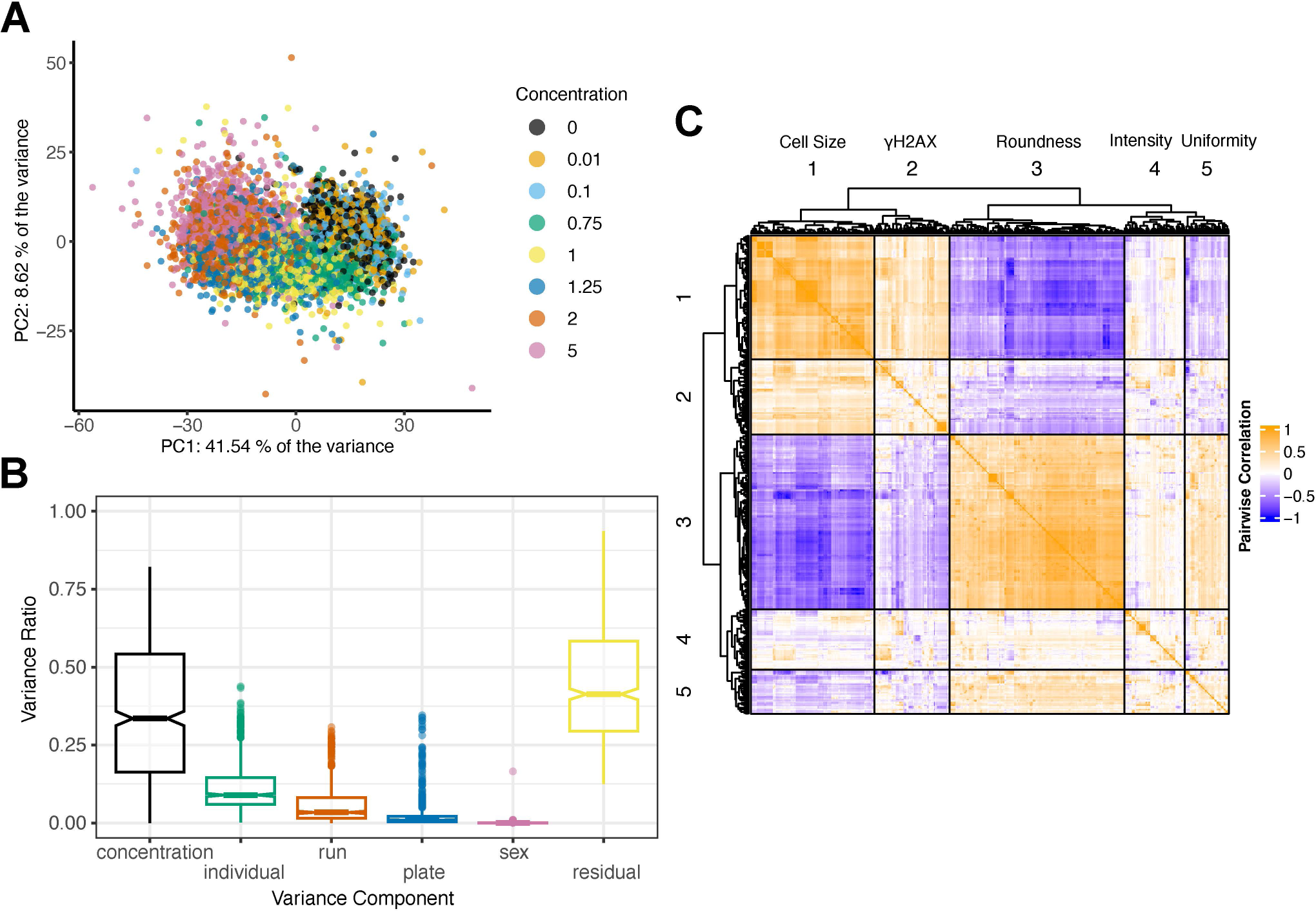
HCS Features are Influenced by MMA^III^ Concentration and Genetic Background. (A) Principal Component Analysis (PCA) of the raw image analysis feature dataset colored by the concentration of MMAIII. Among known factors, increasing MMA^III^ contributed the majority of the variance for both PC1 (41.54 %) and PC2 (8.62%). (B) Boxplot showing the aggregated results from variance component analysis (VCA) performed across all cellular features including MMAIII concentrarions (concentration), DO cell lines (individual), each 96-well plate (plate), residual variation, run, and sex. (D) Heatmap showing the Pearson’s pairwise correlation structure of the all the raw cellular features. The heatmap and dendrogram were generated using the R package ComplexHeatmap’s Heatmap() function with column_split and row_split each set to 5.

While HCS produces thousands of morphological traits, many of them are highly correlated (**Fig. 2C**). The correlated groups could be loosely categorized as traits describing ‘cell sizè, ‘γH2AX focì, ‘cell roundness’, ‘intensity’, and ‘uniformity’ (**Fig. 2C**). While there are a variety of dimension reduction techniques that take advantage of correlation to summarize high dimensional data, we were most interested in traits exhibiting non-linear, dose-dependent responses.

### Dose-response modeling and genetic mapping of cell morphology quantitative trait loci (cmQTL)

Dose-response models are used to define the xenobiotic response profiles of toxicants and drugs. In chemical risk assessment, these models provide benchmark dose estimates, which are the concentrations at which a chemical exposure could pose a health risk ^51^. To focus on the subset of traits exhibiting dose-dependent responses, we performed dose-response modeling using the *drc* R package ^52^ for each cellular trait, individual, and replicate experiment. These models provided quantitative dose-response parameters (DRPs) describing each donor individual’s cellular response including effective concentrations (EC’s), starting/maximum asymptotes, and rates of change (slopes) ^53^. For example, an individual’s EC50 represents the concentration of MMA^III^ at which there is a 50% change in a given cellular feature relative to baseline. Following the removal of redundant features and batch effect correction, our dose-response modeling resulted in 5,105 cmDRPs from 568 cellular traits.

To reveal genetic loci that influence sensitivity to arsenic metabolite MMA^III^, we performed quantitative trait loci (QTL) mapping, treating the 5105 cmDRPs as traits (see Methods). To account for the data’s complicated structure and redundancies in the context of multiple testing burden, we calculated a genome-wide false discovery rate (FDR) significance threshold, which resulted in only the maximum peak meeting significance (FDR < 10%) (**Fig. 3**). Given that this work represents a proof of principle and cmDRPs are potentially noisy as modeled quantities, we also used a lenient significance threshold of LOD score > 7.5, which corresponds to ∼80% genome-wide significance threshold in the DO ^54^. Of the 5105 cmDRPs, 854 possessed suggestive genetic loci associations, with the strongest LOD score being 10.95. We found cmQTL reaching significance on chromosomes 2, 3, 6, 12, 14, 18. Significant response cmQTL included EC’s, slope, and maximum asymptotes, in addition to baseline DRPs, or starting asymptote.

**Figure 3:**
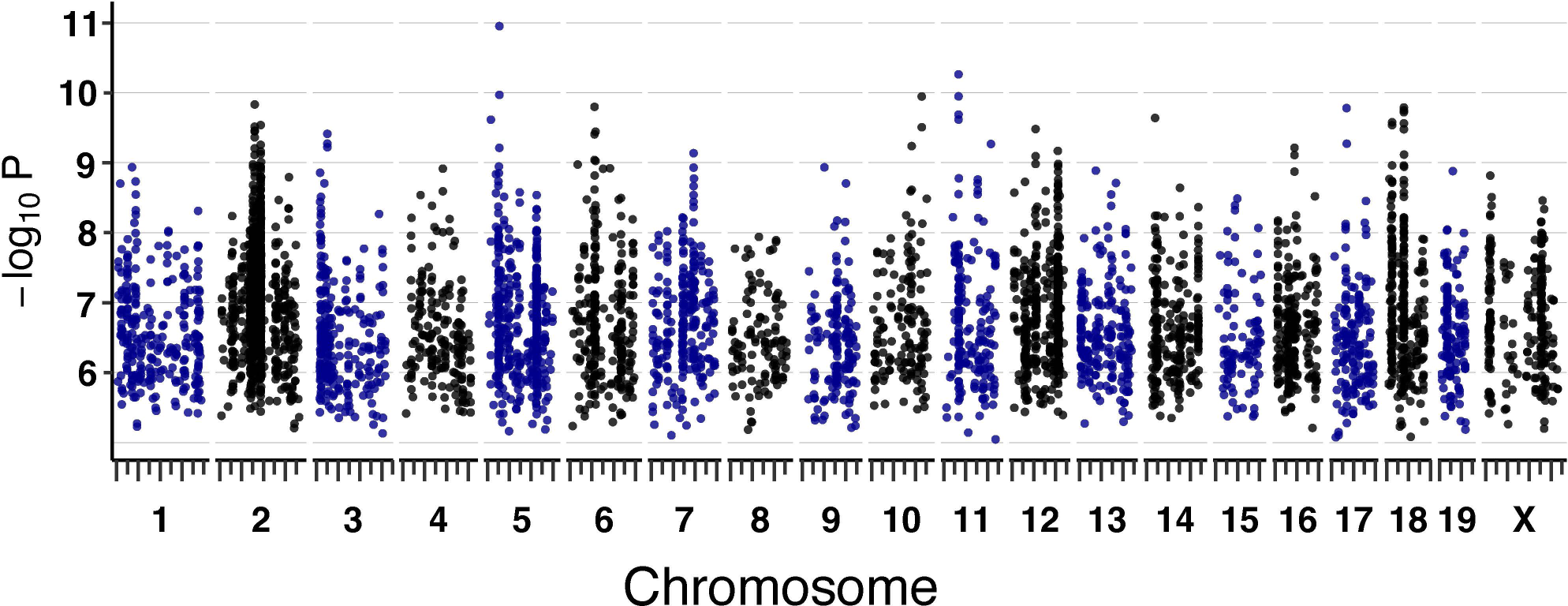
Dose-Response Modeled cmQTL in DO Fibroblasts Exposed to MMA^III^. Summary of cmQTL maximum peaks for 5100 cmDRPs. Each points represents the strength of the genetic association as a LOD score on the y-axis (-log_10_P) across the mouse genome (x-axis). On the x-axis, long tick marks represent the start of the chromosome and 50 Mbp intervals, while the short tick marks are 25 Mbps.

### Candidate cmQTL genes identified using differential gene expression, gene set enrichment, and data integration

To nominate candidate genes and variants within cmQTL, we used several approaches. We generated bulk RNA-Seq data from 16 randomly selected DO fibroblast lines and we used differential expression analysis (DE) to identify expressed genes that showed differential expression in the context of MMA^III^ exposure (**Supp. Table 2**). Then, on the resulting set of genes, we used gene set enrichment analysis (GSEA) to identify groups of genes that are functionally related (**Supp. Table 3**). We interrogated published gene-arsenic interactions through the Comparative Toxicogenomics Database (CTD) ^55^ and for each DE gene, we quantified the number of interaction annotations in CTD across all curated studies involving MMA^III^, MMA^V^, DMA^III^, DMA^V^, sodium arsenite, sodium arsenate, arsenic, and arsenic trioxide. For any causal variants that exert their effects through gene expression, the contributing haplotypes and direction of their effects will be correlated across eQTL and cmQTL in datasets generated from the same genetic reference population (DO). Therefore, we also correlated the cmQTL allele effects with previous DO eQTL from liver, heart, kidney, striatum, pancreatic islet cells, and mESCs (see Methods). Finally, local SNP association mapping within each cmQTL allowed us to identify the SNPs with the highest LOD scores in each interval.

At the pathway level, the most upregulated gene set in dosed samples was ‘NRF2 activation (WP2884)’, which is a well-established response to oxidative stress following arsenical exposure ^56–59^ (**Fig. 4A**). NRF2, also known as NFE2L2, is a transcription factor that is shuttled to the nucleus following dissociation from KEAP1 in response to the generation of ROS ^60–62^. In the nucleus, NFE2L2 binds antioxidant response elements (AREs) upstream of many redox homeostasis and cellular defense genes to drive their transcription in response to stress, including arsenical exposure ^56,57,63–66^. These data provided multiple lines of evidence supporting *Nfe2l2* (*Nrf2*) as a candidate gene for the cmQTL hotspot that we found on Chr 2 (**Fig 3**). Our gene expression analysis also revealed five candidate genes for other response cmQTL with LOD scores > 8 (**Fig. 4B**). Three of the five genes were present within the same CI, including *Hspa1b*, *Hspa1a,* and *Msh5*, with the former two DEGs having over 80 previously defined interactions with arsenicals. Among the other differentially expressed genes we found that 73 (89%) have not previously been associated with MMA^III^, though many have been associated with arsenic or other arsenic metabolites.

**Figure 4:**
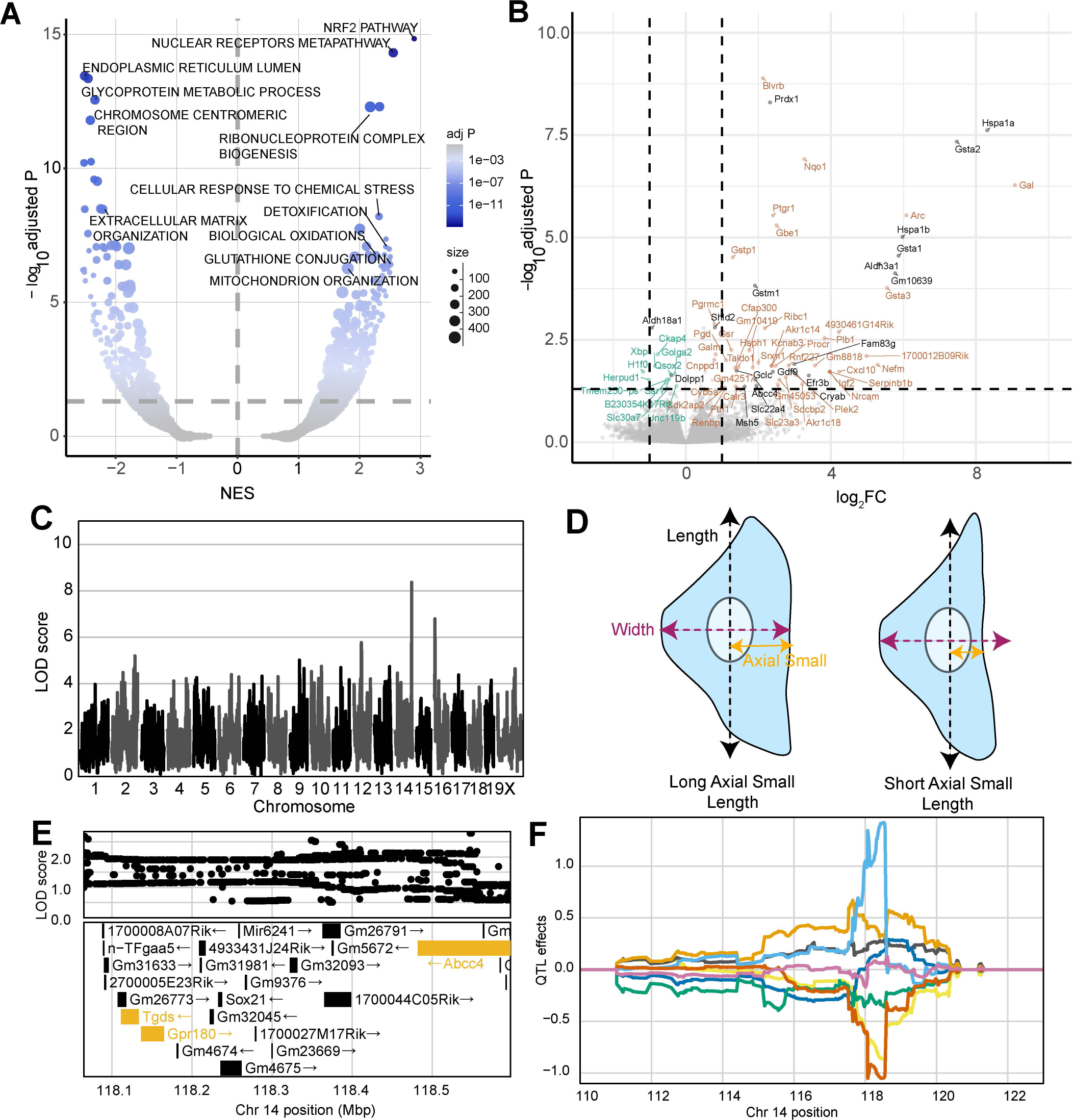
Differential Expression and cmQTL Together Support MMA^III^ Glutathione Conjugation and its Export via ABCC4. (A) Volcano plot showing the normalized effect sizes (NES) and adjusted p-values (-log_10_ transformed) of the score-based gene set enrichment (GSEA) results from differential expression (DE) analysis across the 0 and 0.75 µM MMA^III^ exposed DO fibroblasts groups (n = 32, 16 individuals). Expression was filtered based on a median transcript per million > .5 or removed if at least half of the points were below this cutoff. Each point represent a gene set from ‘GO:Component’,’REACTOME’, ‘KEGG’, ‘WikiPathways’, ‘GO;Tissue’, ‘GO;Molecular Function’ and ‘GO;Biological Process’. The size of each points represents the number of genes within the gene set and the color represents the -log_10_(adjusted P) (y-axis). Horizontal dashed line indicates the adj. p-value significance threshold (adj. P = 0.05) (B) Volcano plot showing the log_2_-fold change (log_2_FC) and adjusted p-values (-log_10_adjusted P) for single genes. The horizontal indicates ithe adj. p-value significance threshold (adj. P = 0.05) and the vertical lines represent the + 1 log_2_fold change for a point of reference. Points labeled with gene names are significantly differentially expressed (adj. p-value < .05) with effect sizes > 0.75 log_2_FC or < -0.25 log_2_FC. Colors represent genes withing cmQTL confidence intervals (black), upregulated (orange) and downregulated (green) DE. (C) QTLscan for the ΈC5 Mitosmooth Axial Small length mean per well’ cmQTL with the maximum peak at chromosome 14: 118483436 bp (m38) and a LOD score of 8.36. (D) Cartoon fibroblast cells depicting the two measurements of cell length (black), width (purple), and axial small width (yellow). Fibroblast on the left has a longer axial small length compared to the fibroblast on the right, (E) Variant association mapping within the Cl the cmQTL ΈC5 Mitosmooth Axial Small length mean per well’. Top panel shows the LOD scores of the known, segregating variants in the 8 DO founders (m38). Bottom panel shows the gene models within the respective Cl. Each point represents a variant. Colors indicate whether a gene is expressed > 0.5 TPM (gold) or < 0.5 TPM (black). The arrow indicates the direction of transcription. (F) Allele effects plot showing the eight DO founders (colors, see Methods) for the ΈC5 Mitosmooth Axial Small length mean per well’ cmQTL across the surrounding region on chromosome 14 (Mbp).

### Natural variation in cellular detoxification pathways partially explains arsenic sensitivity

The other two DEGs within response cmQTL were *Cryab* and *Abcc4,* each with ≥ 19 published arsenical interactions (**Fig. 4B**). SNPs in *Abcc4* have been previously associated with sensitivity to arsenic ^67^. *Abcc4* encodes the protein ABCC4/MRP4, which has been shown to export glutathionylated MMA^III^ from cells ^68,69^. Glutathione transferases like *Gstm1*, *Gsta1,* and *Gstp1* were also significantly upregulated in our expression dataset. These genes are members of the glutathione conjugation pathway which is a detoxification pathway that leads to glutathionylation of MMA^III^ (MMADG^III^) (**Fig. 4A,4B**) ^68,70^. We found multiple cmQTLs at the *Abcc4* locus and they were all for traits related to changes in cell size (i.e., length, compactness) (**Fig. 4C**). For example, one of these response cmQTL was EC5 of the change in axial small length or the dose at which 5% of the cell population exhibited measurable differences in cell size (defined by the smoothed MitoTracker labeling which captures the cytoplasmic area occupied by mitochondria) (**Fig. 4D**). Variant association mapping revealed that the highest scoring SNPs in these cmQTLs were within the *Abcc4* gene, and the allele effects indicated that changes in cell size (‘shrinkage’) occur at lower doses in individuals with PWK haplotypes compared to those with NZO haplotypes (**Fig. 4E,4F**). Taken together, these data support a model where sensitivity to arsenic exposure in the DO population is partly regulated by natural variation in the efficiency of MMA^III^ detoxification.

### *Xrcc2* haplotypes modulate and predict of cellular responses

The cmQTL with the highest LOD score was on chromosome 5 at 27,327,254 bp (GRCm38) for the response cmQTL ‘EC90 Nonborder Nucleus Symmetry 02 SER Hole (Hoechst) Mean Per Well’ (**Fig. 5A, 5D**). Hoechst nuclear fluorescence in cells with the 129 haplotype resembled apoptotic nuclei ^71^ and were brighter and more uniform than those found in cells with AJ/B6 haplotypes (**Fig. 5B, Fig. S1A**). The highest associated SNPs for this cmQTL were located in two genes: *Actr3b* and *Xrcc2* (**Fig. S1B**), however several key points suggest *Xrcc2* as the more likely candidate. First, *Xrcc2’s* paralogs, *Xrcc1* ^72,73^ and *Xrcc3* ^74,75^ have both been associated with genetic susceptibility to arsenical exposure. Second, knockdowns of *Xrcc2* were previously shown to increase both γH2AX intensity and chromosomal abnormalities ^76^, and *Xrcc2* is a member of the Biological Fibroblast Apoptosis (GO:0044346) and DNA Damage Repair pathways (R-MMU-5693532). Lastly, the cmQTL allele effects are highly correlated with an *Xrcc2* eQTL in pancreatic islets cells from the same mouse population (**Fig. 5C**). Taken together, these results suggested that genetic variation at this locus may be mediating DNA damage-induced apoptosis through *Xrcc2* expression.

**Figure 5:**
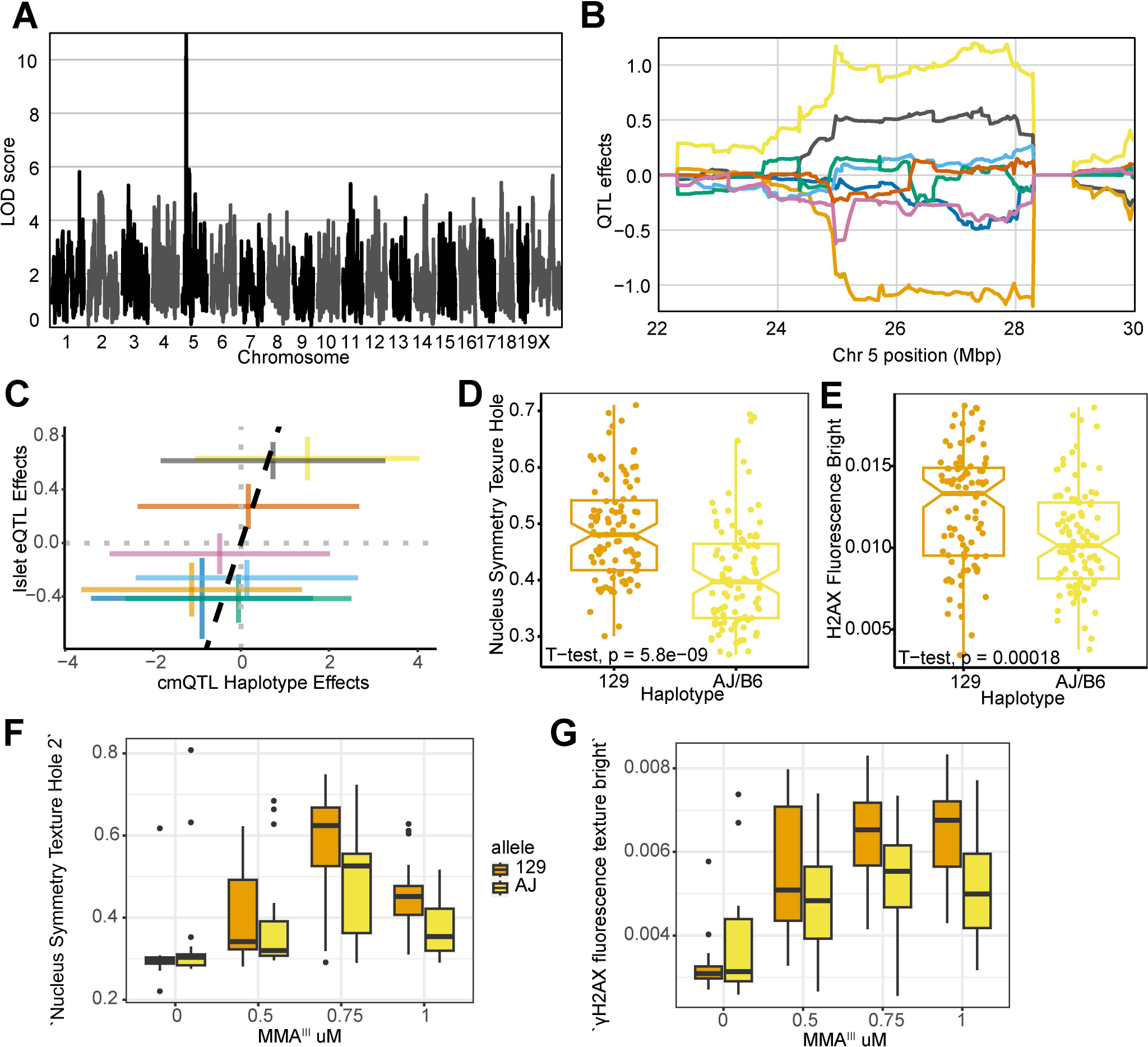
Xrcc2 haplotype modulates chromosomal organization and DNA damage during acute MMA^III^ exposure. (A) QTL scan for the ΈC90 Hoechst Nucleus Symmetry (02) Hole Mean per Well’ cmQTL with the maximum peak at chromosome 5: 27327254 bp (m38) and a LOD score of 10.95. (B) Allele effects plot showing the eight DO founders (colors, see Methods) for the ΈC90 Hoechst Nucleus Symmetry (Hoechst) Hole Mean per Well’ cmQTL across the surrounding region on chromosome 5 (Mbp). Colors indicate founder mouse strains: A/J (yellow), C57BL/6J (gray), 129S1/SvlmJ (orange), NOD/ShiLtJ (dark blue), NZO/HILtJ (light blue), CAST/EĲ (green), PWK/PhJ (red), and WSB/EiJ (purple) (C) Pairwise correlation of the haplotype effects of Xrcc2 expression in pancreatic islet cells at chromsome 5:27,327,254 bp (GRCm38) compared to the haplotype effects of ΈC90 Hoechst Nucleus Symmetry (Hoechst) Hole Mean per Well’. Colors are the same as panel B. (D) Boxplot showing the significant difference (t-test, p value = 5.8e-9) in ‘Nucleus Symmetry Texture Hole 2’ at 1 µM MMA^III^ for the top 129 (n = 24; orange) and AJ/B6 (n = 24; yellow) haplotypes in the DO fibroblasts. (E) Boxplot showing the significant difference (t-test, p value = .00018) in ‘γH2AX fluorescence texture bright’ at 1 µM MMA^III^ for the top 129 (n = 24) and AJ/B6 (n = 24) haplotypes in the DO fibroblasts. (F) Boxplot showing the ΈC90 Hoechst Nucleus Symmetry (02) Hole Mean per Well’ cellular phenotype in a follow-up experiment where DO fibroblasts with 129 (n = 5; orange) and AJ (n = 5; yellow) haplotypes exposed to increasing MMA^III^ concentrations. (G) Boxplot showing the ‘γH2AX fluorescence texture bright’ cellular phenotype in a follow-up experiment where DO fibroblasts with 129 (n = 5) and AJ (n = 5) haplotypes exposed to increasing MMA^III^ concentrations. Colors indicate the DO founder strains (see Methods).

Because of the role in *Xrcc2* in DNA damage and apoptosis, we reasoned that γH2AX fluorescence might also be higher in cells with the more sensitive 129 haplotype compared to cells with the more resistant AJ/B6 haplotypes. Indeed, the γH2AX texture ‘bright’ feature was significantly higher in the fibroblasts with the 129 haplotype compared to the AJ/B6 haplotypes (**Fig. 5E, Fig. S1A**). We sought to assess the reproducibility of these effects, both for the original phenotype and the increase in γH2AX. Taking advantage of our full panel of 600 cell lines, we selected an orthoganal group of lines based on their haplotype at this locus (n = 5 for each allele). Not only were we able to recreate the original nuclear symmetry difference between genetic backgrounds (**Fig. 5F**), but we also observed the same γH2AX fluorescence effects that were found in the original screen (**Fig. 5G**). This example shows that genetic variation in *Xrcc2* influences sensitivity and that the haplotype effects of cmQTL have predictive value for identifying sensitive individuals.

### Non-coding genetic variation influences TXNRD1 cell fate during induced oxidative stress

To further investigate how these data could be used for G x E discovery, cmQTL mapping was performed in a subset of cells lacking accumulated DNA damage. Linear classification was performed to separate cells into H2AX positive and negative populations prior to feature extraction. To do this we took advantage of PHENOLogic machine learning algorithms of the Harmony 4.9 software and gated the imaged cells into γH2AX-negative and γH2AX-positive populations prior to feature extraction, dose-response modeling, and mapping. We detected a cmQTL for the rate of MitoTracker area change in γH2AX-negative cells with a LOD score of 9.16 on chromosome 10 (**Fig. 6A**). This locus was also detected in our original dataset with similar allele effects but with a sub-threshold LOD score (**Fig. S2A, S2B, S2C**). Upon variant association mapping the highest LOD scoring variants were in the 3’-UTR of the *Txnrd1* gene (**Fig. 6C**), a gene that is highly expressed in fibroblasts and has been previously shown to respond to arsenical exposure via changes in NRF2-mediated expression. Moreover, the reducing capacity of TXNRD1 protein is directly inhibited by MMA^III^ binding ^77,78^. As a selenoprotein, the 3’-UTR of *Txnrd1* plays a crucial role in recoding a UGA stop codon into a selenocysteine amino acid which is required for function of the TXNRD1 protein as a reducing agent^79–81^.

**Figure 6:**
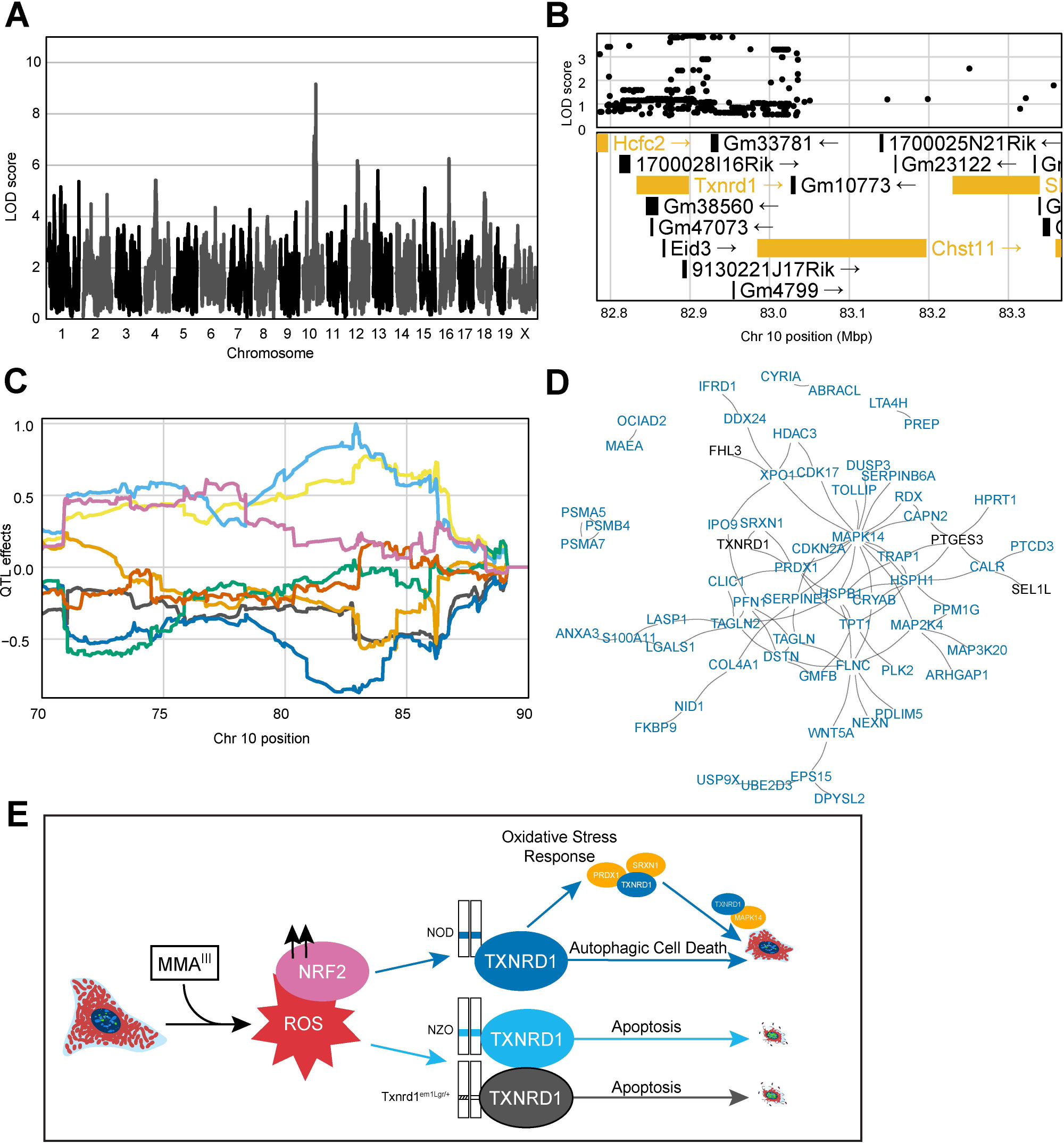
Noncoding Variation in Txnrdl Modulates MMA^III^-lnduced Cell Death. (A) QTL scan for the Ή2AX-negative cells slope Cell Area µm^2^ mean per well’ cmQTL with the maximum peak at Chromosome 10: 82906780 bp (m38) and a LOD score of 9.16. (B) Variant association mapping within the Cl the cmQTL Ή2AX-negative cells slope Cell Area µm^2^ mean per well’. Top panel shows the LOD scores of the known, segregating variants in the 8 DO founders (GRCm38). Bottom panel shows the gene models within the respective Cl. Each point represents a variant. Colors indicate whether a gene is expressed > 0.5 TPM (gold) or < 0.5 TPM (black). The arrow indicates the direction of transcription. (C) Allele effects plot showing the eight DO founders (colors, see Methods) for the Ή2AX-negative cells slope Cell Area µm^2^ mean per well’ cmQTL across the surrounding region on chromosome 10 (Mbp). (D) String-db functional enrichment network of the significantly increased protein interactors detected using immunoprecipitation mass spectrometry (IP-MS) in DO fibroblasts with NOD alleles (n = 6) at the maximum locus for the Ή2AX-negative cells slope Cell Area µm^2^ mean per well’ cmQTL exposed to 0 and 0.75 µM MMA^III^ concentrations. Colors indicate whether a protein, or node, was shared with a similar experiment in DO fibroblasts with the NZO allele (n= 5). Black represents shared TXNRD1 interactors, and blue represents unique NOD-TXNRD1 interactors. (E) Mechanistic summary of allele-specific Txnrdl responses across the NOD haplotype (blue), NZO haplotype (light blue), and heterozygous SECIS knockout model (Txnrdl^em1Lgr/+^). Our data suggest DO fibroblasts with the NOD allele have a more robust oxidative stress response upon MMA^III^ exposure, ultimately succumbing to autophagic cell death represented by increased cell size at medium MMA^III^ concentrations. In comparison, DO fibroblasts with the NZO allele or the Sec^+/-^ alleles undergo a more apoptotic cell fate as shown by brighter Hoechst 33342 labeling and smaller cells.

To interrogate the plausibility of *Txnrd1* as the candidate for these two cmQTL, we performed score-based GSEA using gene expression data from cell lines selected from our collection of 600 lines on the basis of their sensitive (NZO) and resistant (NOD) haplotypes at this locus. We found upregulation of DNA damage and replicative stress gene sets in cells with NZO haplotypes and upregulation of oxidative stress response, p38/MAPK signaling, TGF signaling, RAS signaling, lysosome, and autophagy-related pathways in cells with NOD haplotypes (**Supp. Table 4**). Among these pathways was nanoparticle triggered autophagic cell death, which can be induced by the treatment of gold, the active component of the TXNRD1 inhibitor auranophin ^82^. While we didn’t detect a significant difference in *Txnrd1* transcript abundance by haplotype, at either concentration (**Supp. Table 5**), there was a significant difference in protein levels in the unexposed cells (**Fig. S2D**), and, as expected, TXNRD1 protein levels increased in all arsenic exposed cells. To assess whether TXNRD1 had haplotype specific protein interactions, we performed immunoprecipitation followed by tandem mass spectrometry (IP-MS). Following subtraction of a non-specific binding partner control, we found that compared to healthy, unexposed controls, 0.75 µM MMA^III^ exposed NOD haplotype cells (n = 6) had a larger number (106) of significant, positive interactors compared to NZO (n=5) TXNRD1 interactors (33). NOD TXNRD1 interacted with proteins involved in oxidative stress (i.e., PRDX1, SRXN1), autophagy/p38 (i.e., MAPK14, TOLLIP), and TP53 related REACTOME pathways, while the NZO TXNRD1 interactors did not show pathway enrichment (**Fig. 6D, Supp. Table 6**). Considering the gene expression and IP-MS data together, it was evident that in exposed DO fibroblasts, NOD TXNRD1 was involved in autophagy while NZO TXNRD1 was associated with apoptosis. Previous studies of *Txnrd1* deficiency have shown disruption of lysosomal-autophagy in favor of apoptotic cell death ^83,84^, implying that the apoptotic phenotype of cells with NZO haplotypes (NZO-TXNRD1) is akin to that seen with TXNRD1 deficiency. During apoptotic cell death, cell structure and cytoskeleton are quickly degraded, but during autophagy the cytoskeleton is maintained ^85–87^; providing a basis for our ability to distinguish between these two pathways and to interrogate their genetic regulation using cmQTL. Taken together, these data support a model whereby natural variation in *Txnrd1* influences the trajectory of cell death pathways following MMA^III^ exposure in the DO population (**Fig. 6E**).

While we did not find coding variants unique to the NZO or NOD *Txnrd1* gene, we found that two SNPs private to the NZO haplotype (*rs227869362* and *rs257393906*) in the 3’-UTR were adjacent to the selenocysteine insertion element (SECIS), which is essential for *Sec* recoding during translation. We also searched publicly available data for structural variants and INDELs in the 3’ UTR but did not find any that were unique to the NZO haplotype ^88^. To determine the essentiality of this element *in vivo*, we used CRISPR/cas9 to delete the SECIS in C57BL/6J mice (*Txnrd1^em1Lgr^*). While heterozygous mice carrying this deletion were viable and fertile, homozygous mice could not be recovered. Since a full protein knockout of *Txndr1* causes recessive embryonic lethality ^89^, we concluded that deletion of the SECIS element alone is the functional equivalent of a null allele (see Methods). We then isolated tail tip fibroblasts from heterozygous mice and found that the cell area of arsenic exposed *Txnrd1^em1Lgr/+^* fibroblasts more closely resembled fibroblasts with the NZO haplotype than their WT controls (**Fig. S2E, S2F**). Similarly, nuclear Hoechst 33342 labeling was brighter and more uniform in the *Txnrd1^em1Lgr/+^*nuclei with increasing MMA^III^ concentration. Taken together, these data highlight the functional importance of non-coding variation in the 3’ UTR of a key selenoprotein in the context of sensitivity to arsenic induced oxidative stress. Detailed molecular and functional studies are needed to determine the impact of single nucleotide variants on sec recoding in *Txnrd1*. However, there is at least one study demonstrating that naturally occurring and engineered single nucleotide variants in the 3’ UTR of the human selenoprotein, SEP15, influence UGA readthrough and dampen the cellular response to selenium stimulation ^90^.

### Natural genetic variation influences fibroblast morphology

While our primary focus was on population variation in arsenic response, we unexpectedly observed variation in fibroblast morphology in unexposed cells and our genetic analysis revealed multiple loci contributing to this baseline morphological variation (i.e. starting asymptote cmQTL). The highest scoring of these baseline cmQTL (LOD 9.64) was on proximal chromosome 14 (**Fig. 7A**, **Fig. 7b**). Several of the top LOD scoring variants were in *Ube2e2,* which was one of only three protein coding genes expressed in fibroblasts within the confidence interval (**Fig. 7C**). This cmQTL is for a trait that describes the brightness of Hoechst labeling (i.e., texture feature bright 1 pixel mean per well) which is directly related to the distribution and amount of chromatin in the nucleus (**Fig. 7D**) ^91^. The ubiquitin conjugating enzyme E2 (UBE2E2) functions in the nucleus to post-translationally modify proteins that regulate the G1/S phase transition together with *Trim28* ^92^, which could explain the difference in Hoechst labeling as mitotic cells accumulate more Hoechst due to their DNA content. This example highlights the role of genetic variation in the regulation of morphology, potentially through variation in basic cellular functions (i.e. cell cycle) providing an exciting avenue for further study.

**Figure 7:**
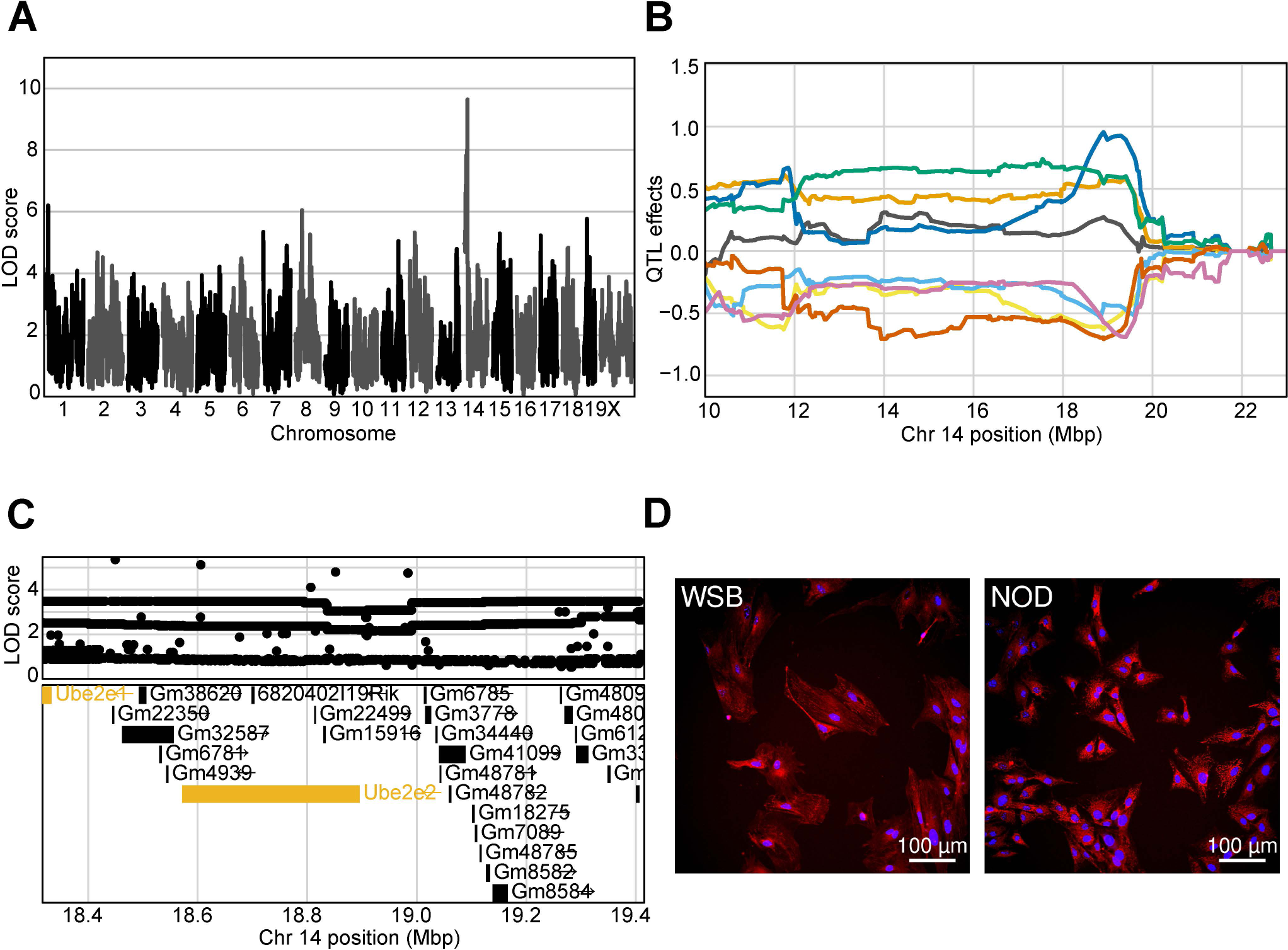
Genetic variation influences fibroblast morphology at baseline. (A) QTL scan for the ‘Hoechst 33342 texture bright 1 pixel mean per well’ cmQTL with the maximum peak at chromosome 14: 19401644 bp (GRCm38) and a LOD score of 9.64. (B) Allele effects plot showing the eight DO founders (colors, see Methods) for the ‘Hoechst 33342 texture bright 1 pixel mean per well’ cmQTL across the surrounding region on chromosome 14 (Mbp). (C) Variant association mapping within the Cl the cmQTL ‘Hoechst 33342 texture bright 1 pixel mean per well’. Top panel shows the LOD scores of the known, segregating variants in the 8 DO founders (m38). Bottom panel shows the gene models within the respective Cl. Each point represents a variant. Colors indicate whether a gene is expressed > 0.5 TPM (gold) or < 0.5 TPM (black). The arrow indicates the direction of transcription. (D) Representative images for the two fibroblast lines showing higher Hoechst 33342 texture bright in the sample with the NOD allele at the chromsome 14 locus comapred to the WSB. Nuclei are labeled in blue by Hoechst 33342 labeling and mitochondria are labeled in red by MitoTracker Deep Red. Scale bar indicates 100 µm.

## Discussion

Taking advantage of a laboratory mouse genetic reference population, we created a new population-based cellular model for *in vitro* analysis of gene by environment interactions. Using this model, we performed HCS to quantify morphological cellular features associated with acute MMA^III^ exposure. We found quantitative variation in these traits across the cell population, and we also found significant variation in the degree to which genetic background could be attributed to this variation (0-40%). We also found significant unexplained residual variation, although the proportion of this contributor to overall variation also varied substantially by trait. Previous studies of cell morphology in genetically diverse cell populations have shown that some traits are prone to high measurement error or experimental variability, especially for features that have high cell to cell variability ^1^. Since our features are whole well summaries, cell to cell variability is a major contributor to our observed residual variation. We also found that the features with higher residual variation were enriched for γH2AX features and that the mean variance ratio for these features was high compared to the overall mean (0.65 vs. 0.4). This higher residual variance is likely due to the indirect immunolabeling method used for γH2AX detection, which is a multistep staining method that relies on two antibodies and is known to have more experimental variability than direct organelle probes.

We used dose-response modeling to summarize cell morphology changes to increasing MMA^III^ insult from which we extracted dose-response parameters (DRPs) as latent traits for QTL mapping. However, there are several notable caveats to this approach. First, to induce cell morphology changes that were likely to fit a sigmoidal dose-response curve, we used concentrations of MMA^III^ that are unlikely to be encountered through environmental or occupational exposures. Other studies have shown that cell morphology was impacted following lower concentration, longer exposures of arsenic ^93^. Secondly, covariates or Bayesian regression during dose-response modeling could allow for better handling of batch effects in high-content imaging data, however these options were not available in the commonly used *drc* R package at the time of our study. Lastly, dose-response modeling varies based on the software being used, the model being fit, and as we observed, the genetic background of the samples. Despite these challenges, we identified hundreds of loci where natural genetic variation in the DO founder strains influences the fibroblast responses to MMA^III^ and baseline fibroblast morphology.

One feature of non-molecular QTL is that while they capture variants with a range of molecular effects (transcriptional or post-transcriptional) they lack a genomic reference point. Thus, a QTL can result from coding variants, noncoding variants, or a combination of both which may influence a cellular trait through a single gene, or multiple, within a QTL region. To refine our cmQTLs and identify candidates, we integrated variant association analysis with orthogonal datasets including gene expression, molecular QTL data from previous DO studies, pathway information, and gene-chemical interaction data from arsenicals through CTD (ctdbase.org). Based on gene-arsenical interactions, we identified 88 genes in our cmQTL that were previously associated susceptibility to arsenic (https://ctdbase.org/). Six genes within our cmQTL including *Abcc4, Nfe2l2, Cbs, Gclc, Gstm1,* and *Xpc* contain SNPs affecting the response to As (https://ctdbase.org/). *Abcc4* was among the significantly differentially expressed genes fibroblasts which make it an intriguing candidate for the EC5 of the change in axial small length cmQTL. Variants in the 3’ UTR of *Abcc4* can regulate its expression through impacting miRNA binding ^94^. We speculate that unique variants in NZO (*rs240728821*) and PWK (*rs245333533*) may be acting in a similar manner. In addition to *Abcc4,* we found 70 novel gene expression changes based on available MMA^III^ exposure within CTD.

Like *Abcc4, Txnrd1* also has an extensive list of gene-arsenical associations in CTD which provides even greater support for the use of ML during image analysis. The ML-derived cell feature, slope H2AX-negative cell area mm^2^, was further corroborated by its presence in the original dataset and by CRISPR-deleted SECIS element in the 3’ UTR of *Txnrd1* recapitulating the same effect. The essentiality of the SECIS element for sec recoding has been previously demonstrated^81^. Our breeding data further support the essentiality of this element for fetal development and our genetic data show that in the 3’ UTR of *Txnrd1* influences the cell size during acute MMA^III^ exposure. The gene expression differences between haplotypes at this locus showed more pro-cancer signaling including RAS, TGF, and p38/MAPK signaling in the NOD haplotype compared to the NZO. This coincides with protein interaction data showing increased NOD TXNRD1 affinity for MAPK14 and oxidative stress related proteins compared to NZO, which may explain the resistance to MMA^III^-induced morphology changes. *Xrcc2*’s involvement in the DNA damage pathway may also indicate a cancer-related outcome for the highest cmQTL ‘EC90 Nonborder Nucleus Symmetry 02 SER Hole (Hoechst) Mean Per Well’. This cmQTL region shares conserved synteny with a region significantly associated with susceptibility to arsenic-induced skin lesions in a Bangladeshi population ^95^.

Fibroblasts are found in many tissues and are involved in disease progression ^96^. However, the genetic effects in fibroblasts may not recapitulate the same molecular mechanisms of sensitivity and resistance as those found in highly specialized cell types. Primary fibroblast cells are also a limited resource because they will undergo senescence, and they are more difficult to genetically manipulate than pluripotent cells. For these reasons, we have generated induced pluripotent stem cell (iPSCs; n = 284) from this panel for future work. iPSCs also enable differentiation into other cell types, 3-dimensional cell models, organoids, or scaffolded arrays which can be screened across a variety of environmental conditions including other toxicants, drugs, or other culture conditions. It is important to note that while other studies mapping cmQTL were limited by lack of genetic diversity, poor adaptation of some cell types to culture, and the genetic architecture of the population being studied ^2,97^, we also found that beyond large effect QTL, our study was underpowered despite previous examples showing sample sizes in this range for molecular phenotypes can detect strong QTL ^12,54,98^. This is the result of experimental and residual sources of variance as described above, as well as the limited extensibility of standard dose-response models to diverse populations. In conclusion, our study demonstrates that dynamic changes in cell morphology ocurring in a population of exposed, genetically diverse cells exhibit predictable dose response relationships. These relationships display interindivual variation and genetic mapping of these relationships unveils the genetic regulation of the molecular initiating events that occur during an acute exposure. Our findings indicate that these loci and their haplotype effects have predictive value for identifying sensitive and resilient individuals *in vitro*. While further work is needed to explore the applicability of these predictions to *in vivo* responses, leveraging mouse genetic reference populations presents an exciting opportunity for iterative *in vitro* screening and precise *in vivo* testing in matched genetic backgrounds.

## Materials and Methods

### Fibroblast Derivation

Tail biopsies approximately 2-3 mm were harvested in from adult male and female Diversity Outbred (RRID:IMSR_JAX:009376) mice, aged approximately 4-6 weeks, using a procedure approved by The Jackson Laboratory’s Institutional Animal Care and Use Committee. Samples were initially collected into Advanced RPMI 1640 cell culture media supplemented with 1.0 % Penicillin Streptomycin (P/S), 1.0 % Glutamax-I (Glutamax), 1.0 % MEM Non-Essential Amino Acids (NEAAs), 0.0005% 2-mercaptoethanol (BME). Tail tissue was minced using razor blades and digested with media containing collagenase D at a concentration of 2.5 mg/ml on an orbital shaker at 37°C. The digested samples were further minced using micropipettes ranging from p1000 to p200 and dissociated in RPMI 1640 media containing 1.0 % P/S, 1% Glutamax, 1.0 % non-essential amino acids, .0005% BME, and 10% fetal bovine serum (FBS), hereinafter referred to as fibroblast media, for approximately 3-5 days (passage number 0; P0). All passaging was done using a phosphate buffered saline pH 7.2 (1X; PBS) wash and 0.05% Trypsin-EDTA (Trypsin). Individual Diversity Outbred fibroblast samples were expanded to P5 with reserve samples frozen at approximate densities of 3.5 x 10^5^ cells/ml at passage numbers P2, P3, and P5 in freeze media containing RPMI 1640 with 10% dimethyl sulfoxide (DMSO) and 10% FBS. All DO fibroblast samples were transferred to liquid nitrogen holding tanks for long-term storage after 24 – 48 hours at -80C.

DNA was harvested from spleen tissue for each DO mouse and samples were genotyped using the Giga Mouse Universal Genotyping Array (GigaMUGA; ^99^). Haplotypes were reconstructed according to the protocol described previously which uses a hidden Markov model to estimate genotype probabilities at each locus for the population ^100^.

### Sample Preparation

Frozen aliquots of P5 fibroblast lines were thawed in fibroblast media and grown for 48 hours in 60 mm tissue culture-treated plates. Viable cell densities were estimated using Trypan Blue (0.4%; Gibco) and a Nexcelom Cellometer Auto T4 Plus Cell Counter. 100 µl of each fibroblast line was seeded into 4 total columns (4 technical replicates) distributed across two CellCarrier Ultra 96-well black, clear bottom, tissue culture treated microplates (PerkinElmer) using the Integra Assist Plus (Integra Biosciences) at a density of ∼2500 viable cells/well following randomization across columns. After 24 hours, fibroblast media was replaced by monomethylarsonous acid (MMA^III^; Toronto Research Chemicals) containing 100 µL of fibroblast media at concentrations of 0 µM, 0.01 µM, 0.1 µM, 0.75 µM, 1.0 µM, 1.25 µM, 2.0 µM, and 5.0 µM in each row which was randomized across plates.

Following 24-hour exposure, MMA^III^ media was replaced with MitoTracker Deep Red (200 nM; Invitrogen) containing media and incubated at 37°C for 20 minutes in the 96-well plates. Subsequently, cells were fixed on ice using ice-cold 100 % methanol for 10-minutes. Following 3X PBS washing, cells were bathed in a 1.0 % bovine serum albumin (Fraction V) (BSA), 0.1 % Tween solution overnight at 4°C on a shaker. After ∼24 hours, blocking solution was replaced with anti-gamma γH2AX antibody (Abcam, ab11174, 1:2000) in blocking solution and incubated at room temperature for 2 hours on a shaker. Following 3X PBS wash, Alexafluor 488 donkey anti-rabbit secondary antibody (1:2000; Abcam) was added for 1 hour at RT on the shaker. After washing, Hoechst 33342 (1:8000; Abcam), was added to cells and incubated for 10 minutes at RT on the shaker. Plates were subsequently washed, and 100 PBS of media was left in each well for storage at 4°C and imaging.

### Automated Image Acquisition

96-well microplates were imaged confocally using an Operetta CLS or Opera Phenix (**Fig. S2E,F**) equipped with a 20x/1.0 water immersion objective and binning 2. A single z-plane was acquired from 25 contiguous fields per well. Exposure times, focal heights, and excitation power settings for the Operetta CLS screen were: Hoechst 33342 (time: 100 ms, power: 100, height: -5), Alexa 488 (time: 200 ms, power: 100, height: -5), MitoTracker Deep Red (time: 500 ms, power: 100, height: -5). Exposure times, focal heights, and excitation power settings for the *Xrcc2* follow-up experiments were: Hoechst 33342 (time: 300 ms, power: 100, height: -6), Alexa 488 (time: 80 ms, power: 100, height: -6), MitoTracker Deep Red (time: 200 ms, power: 100, height: -6). Lastly, exposure times, focal heights, and excitation power settings for the *Txnrd1* follow-up experiments were: Hoechst 33342 (time: 100 ms, power: 80, height: -10) and MitoTracker Deep Red (time: 40 ms, power: 50, height: -10).

### Image Analysis / Cellular Segmentation

‘Basic’ flatfield corrected images were analyzed and processed using Harmony 4.9 software with PhenoLOGIC (PerkinElmer). Gaussian smoothed images were used for image segmentation, with a focus on 2 main regions of interest (ROIs) including using Hoechst 33342 to define the nucleus, and MitoTracker Deep Red to define the cytoplasm surrounding each nuclear ROI. Fluorescence patterning (i.e. texture) and intensity were measured in the nuclear and cytoplasmic regions using the Hoechst 33342 and MitoTracker Deep Red/MitoTracker Deep Red Gaussian smoothed channels, respectively. Features including nuclear area, Hoechst 33342 intensity, and nucleus edge texture were extracted and represented as mean +/- SD per well.

The second image analysis approach used the PhenoLOGIC machine learning (PerkinElmer) algorithms in the Harmony 4.9 software define sub-populations of cells based on γH2AX/Alexa-488 secondary labeling (γH2AX positive and γH2AX-negative) and MitoTracker Deep Red (stressed and unstressed) prior to feature extraction to generate features including ‘MitoTracker Cell Area in γH2AX negative cells’.

### Feature Variance and Relatedness

Principal components analysis was performed on the image analysis features across all concentrations, individuals, and plates using the ‘pcà function from the R *pcaMethods* with the option ‘scale = “uv”’. Variance component analysis was performed using the ‘lmer’ function from the R package *lme4*. The sources of variation included in the model were sex, DO generation (‘generation’), DO donor (‘individual’), 96-well plate (‘plate’), and run (See **Equation 1)**. Variance components were extracted from the model using the function ‘VarCorr’ for each of the random effect (generation, sex, individual, and plate). Residual variance was extracted as the sigma from the model summaries. Ratios of the variance components were determined by dividing each variance component by the sum of all the variance components and the residual variance.

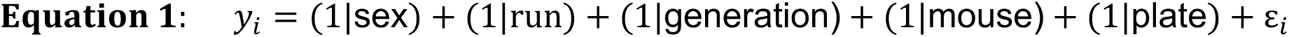

Lastly, the pairwise correlation structure of these data was calculated using the ‘cor’ function in the *WGCNA* R package with the option ‘use = “pairwise.complete.obs”’. The heatmap was created using the *ComplexHeatmap* R package with the dendrogram added using the ‘column_split’ and ‘row_split’ options each set to 5. We added terms to the heatmap clusters based on a qualitative examination of the clustered trait names.

### Cellular Feature Dose-Response Modeling

We used the *drc* R package ^52^ to perform dose-response modeling for each of (insert total number) cellular features. For each of (how many) individuals, we fit 4 technical replicates to the four-parameter log-logistic dose-response model (see Equation 2) using the ‘drm’ function with the ‘fct’ set to ‘LL.4’ and log-normalized cellular features using the ‘bcVal = 0’ option. Model parameters, as shown in Equation 2 ^52^ where *x* represents concentration, including slopes (*b*), upper asymptotes (*d*), lower asymptotes (*c*), and EC50’s (*e*) were extracted from the summary of the model fits. Additionally, the ‘ED’ function was used to estimate the EC5, EC10, EC25, EC75, and EC90 for each model fit ‘relative’ to the asymptotes. 4 replicates for each model fit parameter summary were estimated for each DO individual and cellular feature.

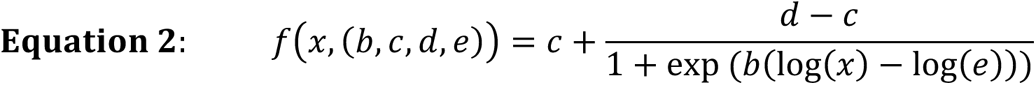

### LMM / BLUP Estimation

Samples were analyzed on different days, across many 96-well plates, and multiple MMA^III^ exposures. We summarized the dose-response parameter replicates using Equation 3 accounting for each individual and plate as random effects. We adjusted for potential batch effects across DO progenitors’ concentration response parameters using linear mixed effect models (LMM). We fit the LMM using the ‘lmer’ function from the R package *lme4*. We modeled each cellular feature as where *y_i_* is the dose-response parameter estimate for a given cellular feature for DO progenitor *i*, modeled with varying intercepts through random effects for mouse/progenitor and 96-well plate ε_*i*_ is the random error term, assumed to ε*_i_* ∼ N(0, σ^2^), and σ^2^ is the error variance. Data without the effect of plate were extracted as the best linear unbiased predictors (BLUPs) of the random effect for DO progenitors and used for QTL mapping analysis.

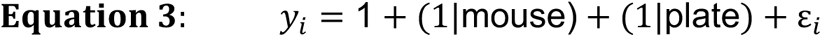

### Cellular Feature QTL Mapping

All data were converted to the normal quantiles calculated from the ranked data, i.e., the rank-based inverse normal transformation (rankZ) to force a Gaussian distribution for mapping. QTL mapping was performed using the *qtl2* R package. Briefly, a genetic relationship matrix (i.e., kinship matrix) was calculated from the genotype probabilities using the ‘calc_kinship’ function with the ‘leave one chromosome out’ (loco) option for genetic mapping and the “overall” option for heritability (*h*^2^) estimation. Sex and DO generation were included as covariates following One hot encoding in the LMM for both heritability estimation and QTL mapping.

For QTL mapping, we first tested individual loci spanning the genome for association with each cellular feature (using qtl2’s ‘scan1’ function). We then estimated allele effects at detected QTL as BLUPs (using the ‘scan1blups’ function) to identify the parental haplotypes driving each QTL and their respective directionality. SNP-association mapping was performed using the ‘scan1snps’ function and the known variants across the eight founder strains of the DO. We calculated a genome-wide false discovery rate (FDR = .10) using the permutations (n = 1000) for the ‘EC50 number of nucleì trait as simulated permutations for all 5105 cmDRPs mapped.

### Diversity Outbred Fibroblast MMA RNA-seq preparation

32 cell lines, including those with NOD or NZO haplotypes at Chr10:82.89 (GRCm38) were thawed into 60 mm cell culture treated plates and grown to confluency (≥ 0.8 x 10^6^ cells/ml) in DO media. Each cell line was then passaged equally into 2 60 mm cell culture dishes and grown to 75% confluency upon which 1 60 mm dish received 0.75 µM MMA^III^ containing DO media and 1 60 mm dish received standard DO media. Following 24-hr exposure, both treated and untreated samples were independently collected and snap frozen on dry ice as cell pellets for 15 minutes. Samples were stored at -80°C prior to RNA isolation. RNA was extracted using a NucleoMag RNA Kit (Macherey Nagel) and purified with a KingFisher Flex system (ThermoFisher). Library preparation was enriched for polyA containing mRNA using the KAPA mRNA HyperPrep Kit (Rocher Sequencing and Life Science). Paired end sequencing was performed with a read-length of 150 bp on an Illumina NovaSeq 6000.

### Transcriptomic Profiling

Genotypes for each sample were then reconstructed using the genotype by RNA-seq pipeline (GBRS) and aligned to the 8 founder allele-specific genome using GBRS RNA-seq pipeline to quantify read counts for each gene ^101^ (available through Github at TheJacksonLaboratory/gbrs_nextflow. These expected counts were the input for differential expression between the 0 and 0.75 µM exposures using the R package *DEseq2* ^102^. We then used the *fgsea* R package to perform a score-based gene set enrichment analysis ^103^. The input for GSEA was the exposure-based log_2_ fold-change for each gene normalized by its standard error. Gene Ontology (GO), REACTOME, WikiPathways, and Biocarta genesets for *mus musculus* were obtained via the R package *msigdb* ^104^. Additionally, the R package *ClusterProfiler* was to assess enrichment of the significant differentially expressed gene set based on the outlying alleles for the cmQTL on chromosome 10 (GRCm38) ^105^.

### CTD Database Mining

The Comparative Toxicogenomics database (CTD) was used to identify gene-arsenic interactions previously defined for candidate genes within cmQTL CIs. The gene-arsenic interactions were downloaded for these arsenicals: monomethylarsonic acid (MMA^V^), monomethylarsonous acid (MMA^III^), dimethylarsinic acid (DMA^V^), dimethylarsinous acid (DMA^III^), arsenic trioxide (ATO), sodium arsenite, sodium arsenate, and elemental arsenic (As). NCBI gene ID’s were then merged to Ensembl IDs and their mouse orthologs obtained through Ensembl’s BioMart tool ^106^. We aggregated the number of ‘Interactions’ for each gene across the arsenicals to get an ‘Interaction Count’ for the genes within cmQTL CIs.

### TXNRD1 Relative Abundance

DO fibroblasts were selected based on their genotypes at the *Txnrd1* locus representing 6 NOD, 5 NZO, and 4 NOD/NZO haplotypes balanced for both male and female lines. Each line was split into two 60 mm dishes where one 60 mm plate received 0 µM MMA^III^ containing media (unexposed) while the other contained 0.75 µM MMA^III^ containing media. After 24 hours, cell pellets split into two vials and snap frozen on dry ice for further processing and liquid chromatography tandem MS (LC-MS/MS) analyses. Protein pellets were resuspended in 150 uL of 50 mM HEPES, pH 7.4, and lysed by passing through a syringe with 28 gauge needle (10 passes), vortexing for 30 seconds, and waterbath sonicating for 5 minutes (30 seconds on, 30 seconds off). Lysates were then clarified via centrifugation at 21,000 x g for 10 minutes at 4°C. Clarified lysates were quantified using a microBCA assay and 20 µg samples were diluted to 50 uL for digestion in 50 mM HEPES, pH 8.2. Samples were then reduced with 10 mM DTT at 37°C for 30 minutes, alkylated with 15 mM IAA at room temperature in the dark for 20 minutes, and trypsin digested overnight at 37°C (trypsin:protein ratio of 1:50). Samples were then cleaned-up using Millipore P10 zip-tips, dried in a vacuum centrifuge, reconstituted in 20 µL of 98% water/2% ACN with 0.1% formic acid, and transferred to mass spec vials. Each sample was analyzed using Thermo Eclipse Tribrid Orbitrap Mass Spectrometer coupled to a nano-flow UltiMate 3000 chromatography system on a Thermo 50 cm EasySpray C18 column as described previously with the exception that the gradient was scaled down to a 90 minute gradient^107^. TXNRD1 abundance was determined based on the target peptide: IEQIEAGTPGR. Raw peak data was processed using Skyline (version 22.2.1.278) and further analyzed in R. Significance across alleles and concentrations was assessed using permutations (n = 1000) because of the non-normal distributions of the protein levels. All mass spectrometry analysis was performed in the in The Jackson Laboraory (JAX) Mass Spectrometry and Protein Chemistry Service.

### Immunoprecipitation Mass Spectrometry (IP-MS)

Immunoprecipitation mass spectrometry (IP-MS) was performed using a rabbit antibody derived against the mouse TXNRD1 protein gifted from Dr. Edward Schmidt to determine TXNRD1 binding partners using the samples and instrumentation described in the ‘TXNRD1 Relative Abundance’ section. M-280 Sheep Anti-Rabbit IgG Dynabeads (Invitrogen, 11203D) were prepared and coupled to the rabbit anti-mouse TXNRD1 antibody according to manufacturer protocol; additional IgG control beads with no TXNRD1 were also prepared as a non-specific binding partner control for the beads. A ratio of 5 ug of antibody to 5 x 10^7^ beads was used. All Dynabeads were then blocked with 5 mg/mL BSA overnight at 4°C during the antibody coupling step. Coupled and control IgG Dynabeads were then bound to 250 µg of protein lysate at room temperature with rotation for one hour. Heterozygous samples were pooled and used as IgG subtractive controls to assess non-specific binding for the beads. All bound bead fractions were clarified with a magnet, then washed three times with Wash Buffer A (10 mM HEPES at pH7.4, 10 mM KCl, 50 mM NaCl, 1 mM MgCl2, NP-40 (0.05% w/v)), followed by two washes with Wash Buffer B (10 mM HEPES at pH7.4, 10 mM KCl, NP-40 (0.05% w/v)). Washed beads were then digested on-bead as described for the relative abundance section above with the exception of 500 ng of trypsin being used. Samples were then purified using a Millipore P10 Zip-tip and prepped for tandem mass spectrometry analysis, both as described above in the relative abundance section. Raw data was analyzed using the Thermo Proteomic Discoverer software as described previously in the JAX Mass Spectrometry and Protein Chemistry Service using standard operating protocols ^107^.

### PPI and Functional Enrichment

We used the *string_db* R package to assess functional enrichment of proteins binding TXNRD1 to generate protein-protein interaction (PPI) networks for the allele-specific IP-MS results ^108^. We used a score threshold of ‘400’ to identify functional interactions between TXNRD1 interacting proteins (nodes) across NOD and NZO haplotypes at the chromosome 10 locus which were indicated as edges in the *igraph* R package visualization. The PPI was colored based on shared (black) and unique (blue) proteins across alleles.

### *Txnrd1* SECIS deletion

To delete a 200 bp domain containing the SECIS regulatory element of *Txnrd1* (MGI:1354175, NCBI Gene: <u>50493</u>, ENSMUSG00000020250) as well as the flanking regions where 3’ UTR variants are found in NZO haplotypes, we engineered C57BL/6J (The Jackson Laboratory stock #000664, RRID:IMSR_JAX:000664RRID:JAX000664) embryos using CRISPR/Cas9. The SECIS element of murine *Txnrd1* is a 75 bp regulatory element ranging from 1967-2042 bp in NM_001042513.1, essential for recoding UGA to specify selenocysteine. Two sets of gRNAs were used (gRNA up 1:GGAGGCTGCAGCATCGCACT, gRNA down 1: GGGTTAATGATACTAGAGAT, gRNA up 2: GAGGCTGCAGCATCGCACTG, gRNA down 2: GGTTAATGATACTAGAGATA) with no repair template. Off-target effects were assessed using the Benchling algorithm (https://benchling.org) and for all guides, potential off target sites were scored <2.0. Two F0 founders (male 5007 and female 5016) carrying the expected 220 bp deletion at chr10:82,896,230-82,896,450 (GRCm38) were identified by PCR. PCR genotyping primers were designed to amplify a 565 bp WT product and a 365 bp deletion product (SECIS_500_FWD 5’ CCTTCCTCTTT CTGCAGATATT 3’, SECIS_500_REV 5’ ACC CAC TTCCACACAGTAAAG 3’). Male founder 5007 was backcrossed to C57BL/6J females and PCR genotyping (primers) was used to identify N1 heterozygous offspring. After two more backcrosses N3 animals were intercrossed to generate N3F1 and N3F2 animals for phenotyping and tail tip fibroblast biopsy. The heterozygous crosses resulted in 320 animals, 211 animals were heterozygous (66%), 109 were wildtype (34%) and 0 were homozygous for the deletion allele. This 2:1 Mendelian ratio (het:WT) was consistent with recessive embryonic lethality of the deletion allele. Targeted oxford nanopore sequencing of the was used to confirm the sequence of the deletion allele and the lack of closely linked off target mutations in the *Txnrd1* gene. The resulting strain C57BL/6J-*Txnrd1^em1Lgr^*/Lgr was assigned The Jackson Laboratory stock #37668. All experiments using mice were approved by The Jackson Laboratory’s Institutional Animal Care and Use Committee.

## Data and Code Availability

All statistical analyses were performed using the R statistical programming language (v4.1.3)^109^. The data, supplemental tables, and analysis pipelines used to process, analyze, report, and visualize these findings are publicly available (10.6084/m9.figshare.24576181). The raw and processed RNA-seq data are available from Gene Expression Omnibus (GEO) (GSE247877). All images are available from the corresponding authors (C.O.,L.R.) upon reasonable request.

## Author Contributions

Conceptualization, L.G.R, R.K., G.C., C.O.; Methodology, L.G.R., C.O., W.M., G.R.K.; Validation, C.O., W.M., G.R.K., L.G.R.; Formal analysis, C.O., G.R.K., D.G., G.C.; Investigation, C.O., W.M., B.R.H., T.S., L.G.R. Resources, R.K., G.C., D.G.; Writing – Original Draft, C.O., L.G.R, G.R.K.; Writing – Review and Editing, R.K., G.C., L.G.R., D.G., B.R.H., G.R.K., C.O.; Visualization, C.O.; Supervision and Project Administration, L.G.R., G.C., R.K.; Funding acquisition, L.G.R., G.C., R.K., D.G.

## Acknowledgements

We thank Drs. Edward Schmidt and Dr. Justin Prigge, Montana State University for providing the TXNRD1 antibody. We also acknowledge the support of The Jackson Laboratory Mass Spectrometry and Protein Chemistry Service, Protein Sciences, The Jackson Laboratory Genome Technologies Core, and Jackson Laboratory Computational Sciences for their expert assistance. We also thank Dr. Belinda Cornes and Robert Sellers for their computational support during the early stages of this project. Lastly, we thank Dr. Stephen Straub at PerkinElmer for his support including reviewing this manuscript.

---

**Figure S1:**
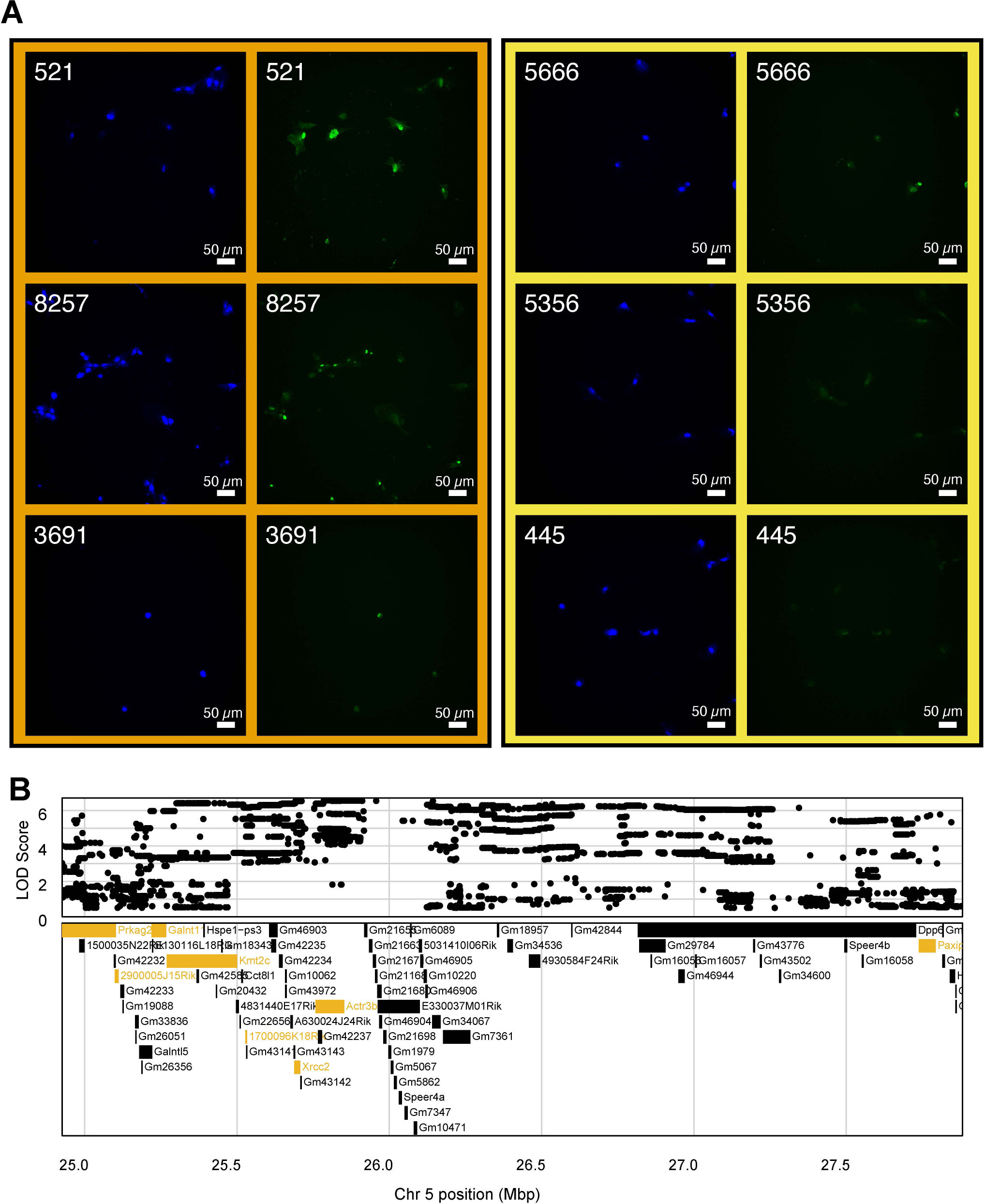
Genetic variation near *Xrcc2* associated with nuclear changes following MMA^III^ exposure. (A) Representative images for the two fibroblast lines at a 1 µM MMA^III^ concentration with nuclei labeled by Hoechst 33342 (blue) and y H2AX (Alexa-488 secondary; green) for primary fibroblasts with a 129 allele (orange; n = 3) versus an AJ/B6 allele (yellow; n = 3) at the maximum position for the ΈC90 Hoechst Nucleus Symmetry (02) Hole Mean per Well’ cmQTL. (B) Variant association mapping within the Cl the cmQTL ΈC90 Hoechst Nucleus Symmetry (02) Hole Mean per Well’. Top panel shows the LOD scores of the known, segregating variants in the 8 DO founders (GRCm38). Bottom panel shows the gene models within the respective Cl. Each point represents a variant. Colors indicate whether a gene is expressed > 0.5 TPM (gold) or < 0.5 TPM (black).

**Figure S2:**
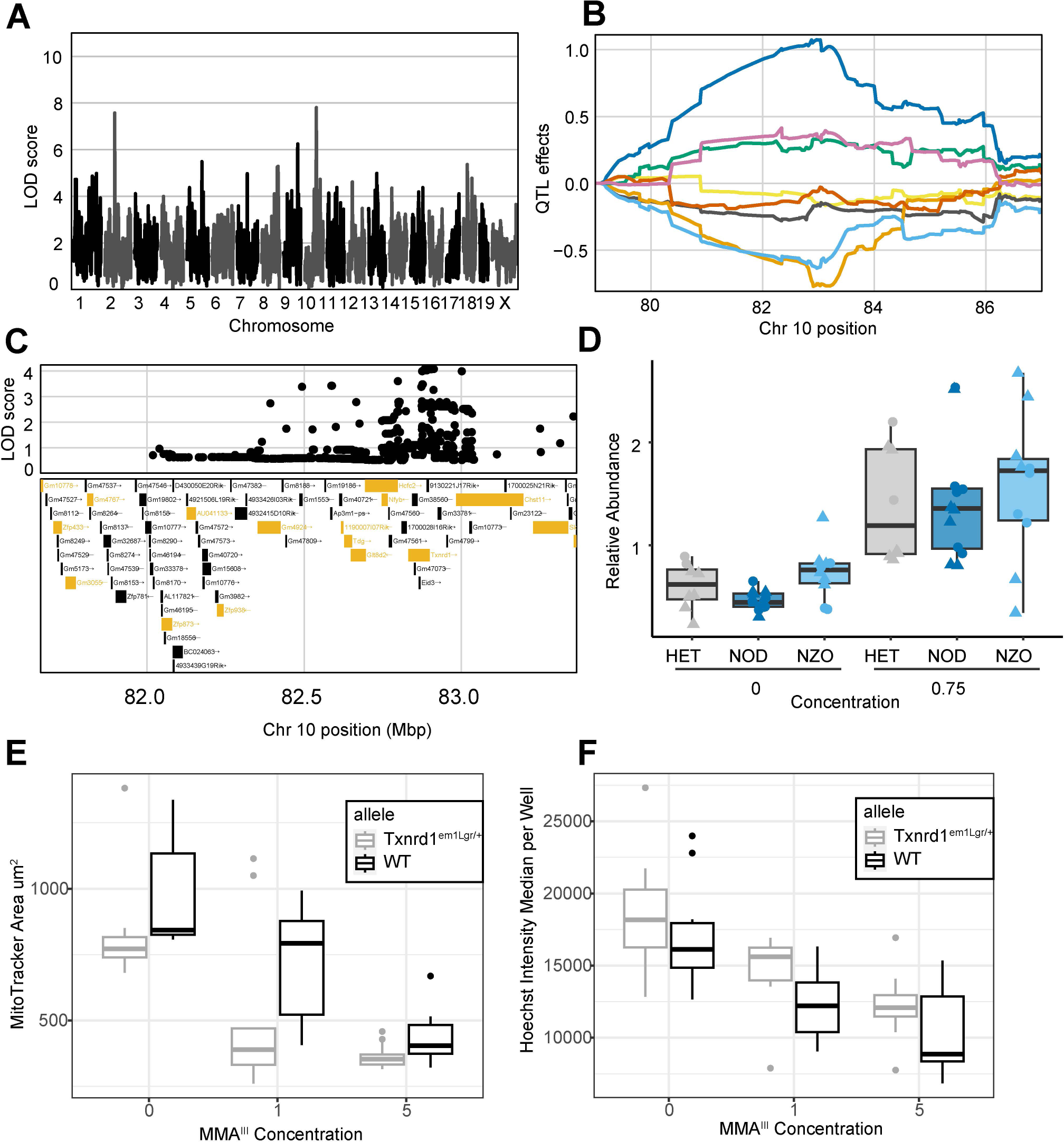
Heterozygous SECIS-Knockout in TxnrcH Recapitulates Cell Area Phenotype. (A) QTL scan for the ΈC90 Mitosmooth Symmetry (3) Texure Edge Mean per Well’ cmQTL with the maximum peak at chromosome 10: 82,967,807 bp (m38) and a LOD score of 7.64. (B) Allele effects plot showing the eight DO founders (colors, see Methods) for the ΈC90 Mitosmooth Symmetry (3) Texure Edge Mean per Well’ cmQTL across the surrounding region on chromosome 10 (Mbp). (C) Variant association mapping within the Cl the cmQTL Ή2AX-negative cells slope Cell Area µm^2^ mean per well’. Top panel shows the LOD scores of the known, segregating variants in the 8 DO founders (m38). Bottom panel shows the gene models within the respective Cl. Each point represents a variant. Colors indicate whether a gene is expressed > 0.5 TPM (gold) or < 0.5 TPM (black). The arrow indicates the direction of transcription. (D) Relative abundance of TXNRD1 compared between DO fibroblast lines with NOD (n=6), NZO (n=5), and NOD/NZO (n=4) alleles at the chromosome 10 locus. Significance testing was performed using permutations testing (n = 1000). Star (*) represents p-value < .05. (E) MitoTracker Deep Red Cell Area across increasing MMA^III^ concentration for Txnrd1^em1Lgr/+^(n=3) compared to B6 control (n=3) primary fibroblasts. Colors indicate wild-type (black) compared to Txnrd1^em1Lgr/+^(gray) primary fibroblast lines. (F) Ήoechst 33342 intensity’ across increasing MMA^III^ concentration for Sec^+/-^(n=3) compared to B6 control (n=3) primary fibroblasts. Colors indicate wild-type (black) compared to Txnrd1^em1Lgr/+^(gray) primary fibroblast lines.

